# Highly contiguous genome assembly and population sequencing reveal three distinct populations and signatures of insecticide resistance of *Tuta absoluta* in Latin America

**DOI:** 10.1101/2022.12.15.520645

**Authors:** Kyle M. Lewald, Christine A. Tabuloc, Kristine E. Godfrey, Judit Arnó, Clérison R. Perini, Jerson C. Guedes, Joanna C. Chiu

## Abstract

*Tuta absoluta* is one of the largest threats to tomato agriculture worldwide. Native to South America, it has rapidly spread throughout Europe, Africa, and Asia over the past two decades. To understand how *T. absoluta* has been so successful and to improve containment strategies, high quality genomic resources and an understanding of population history is critical. Here, we describe a highly contiguous annotated genome assembly, as well as a genome-wide population analysis of samples collected across Latin America. The new genome assembly has an L50 of 17 with only 132 contigs. Based on hundreds of thousands of SNPs, we detect three major population clusters in Latin America with some evidence of admixture along the Andes Mountain range. Based on coalescent simulations, we find these clusters diverged from each other tens of thousands of generations ago prior to domestication of tomatoes. We further identify several genomic loci under selection related to insecticide resistance, immunity, and metabolism. This data will further future research toward genetic control strategies and inform future containment policies.

## Introduction

*Tuta absoluta* (also known as *Phthorimaea absoluta* (Chang & Metz 2021)) is a worldwide economic pest of tomatoes and other solanaceous crops. A member of the gelechiid family, this moth lays eggs on the above-ground portion of the plant, where the hatched larvae will spend their lives creating “mines” throughout the plant tissue before pupating and emerging as adults (Godfrey et al. 2018). At a reproduction rate of up to 10 generations per year, untreated infestations will eventually result in complete death of the plant, leading to up to 100% agricultural loss. Although a large effort has been made to develop and implement integrated pest management (IPM) programs across different world regions (Desneux et al. 2022), typical treatments have included heavy use of a variety of insecticides (Siqueira et al. 2000), leading to the rapid appearance of insecticide resistance. As tomatoes represent a massive economic industry, with an estimated 252 million metric tons of tomatoes harvested in 2020 (FAOSTAT 2020), there is a serious need to understand the invasive biology of this insect and to develop tools for detection and prevention.

*T. absoluta* was originally detected in Peru in 1917 (Meyrick 1917) but was not recorded as an agricultural pest until the 1960s and 70s when it was discovered in tomato fields in Chile, Argentina, and Venezuela; by the 1990s it was widespread across South America. In 2006, *T. absoluta* appeared in Spain (EPPO 2008); since then it has rapidly colonized Europe, Asia, and Africa. It is generally believed that the Peruvian highlands is the ancestral home of *T. absoluta*, and that the rapid colonization to the rest of Latin America was due to the introduction of *T. absoluta* by human transport of contaminated fruit, although few studies have confirmed this (Desneux et al. 2011). Previous research using mitochondrial and microsatellite DNA markers found some evidence of population structure, as well as evidence that the European invasion originated from a single population in central Chile (Cifuentes et al. 2011; Guillemaud et al. 2015). However, determination of higher-resolution population structure, migration events, divergence times, and population size can benefit from using a larger number of markers, such as what is produced from genome-wide sequencing studies (Trask et al. 2011; Willing et al. 2012; Rašić et al. 2014; Koch et al. 2020). Additionally, few genetic studies have been conducted to understand how *T. absoluta* has performed so successfully as an agricultural pest beyond targeted examinations of known insecticide resistance alleles. One reason for this has been the lack of a highly contiguous genome with annotated genes. A short-read based assembly has been previously published for the purpose of developing molecular diagnostics (Tabuloc et al. 2019); however, it is highly fragmented and duplicated.

In this study, we addressed these issues by using long-read sequencing technology to produce a highly contiguous genome assembly for *T. absoluta*. We then use short-read technology to sequence genomes of individuals collected across Latin America, as well as a Spanish population, to identify single nucleotide polymorphisms (SNPs) in an unbiased manner. We use these SNPs to detect population structure and estimate population history parameters to understand how and when *T. absoluta* spread across Latin America. Finally, we use genome scanning statistics to identify genes putatively under selection that may explain *T. absoluta’s* success as an agricultural pest. We expect the genome assembly and population data will be an asset toward developing new strategies to manage this pest.

## Methods

### High Molecular Weight DNA extraction

For genome assembly, a single *T. absoluta* larva was collected from a colony originally sourced from the Institute of Agrifood Research and Technology (IRTA), Cabrils, Spain and held in the Contained Research Facility in UC Davis and frozen on dry ice. The larva was pulverized in liquid nitrogen with a pestle in a 2 mL microcentrifuge tube using 740 mL of lysis buffer (80 mM EDTA pH 8, 324 mM NaCl, 0.68% SDS, 8 mM Tris-HCl pH 8, 80ug/mL RNase A (NEB, Ipswich, MA), 135 ug/mL Prot K(NEB)). After a 37°C overnight incubation step, 240 μL of 5M NaCl was added and gently mixed in by rocking before centrifuging at 10,000 RCF, 4°C, for 15 minutes. Supernatant was transferred using a wide-bore pipette to a 2 mL DNA low-bind tube (Eppendorf, Enfield CT), precipitated with 1 mL of 100% ethanol, and centrifuged at 10,000 RCF, 4°C, for 5 minutes. The DNA pellet was washed with 500 μL of ice-cold 70% ethanol twice before air-drying for 5 minutes. Dry pellet was resuspended in DEPC-treated water and allowed to dissolve for 1 hour at room temperature before being stored at 4°C for no more than 2 weeks. Absorbance ratios were measured with a Nanodrop Lite (ThermoFisher Scientific, Waltham, MA), DNA concentration was measured with a Qubit 4 Fluorometer using a dsDNA High-Sensitivity Assay (ThermoFisher Scientific), and DNA fragment size was measured with a Tapestation genomic DNA ScreenTape (Agilent, Santa Clara, CA). Approximately 700 ng of DNA was sent to QB3-Berkeley for library preparation and PacBio HiFi sequencing with 1 SMRTcell.

### Genome Assembly and Assessment

Raw subreads were collapsed into Circular Consensus Sequence (CCS) reads using ccs version 6.0.0 (PacBio, Menlo Park, CA). K-mer histograms were made with jellyfish version 2.2.6 (Marçais & Kingsford 2011) using 31-mers, then visualized with GenomeScope version 2.0 (Ranallo-Benavidez et al. 2020). GC content vs k-mer frequency was calculated from the jellyfish histograms using kat version 2.4.2 (Mapleson et al. 2016) and visualized with R.

CCS reads were initially assembled using hifiasm version 0.14 or HiCanu version 2.11 (Nurk et al. 2020; Cheng et al. 2021) with default parameters. HiCanu assembly was separated into a primary and alternate haplotig set using purge-dups version 1.2.5 4 (Guan 2021). The primary hifiasm assembly was purged with purge_dups; alternate haplotigs from this purge were purged with the alternate hifiasm assembly, and re-purged with purge_dups to discard repeats, high-coverage, or nested haplotigs.

Merqury version 1.1 (Rhie et al. 2020) was used to assess genome assembly quality and completeness between the two assemblers and between pre- and post-purging. K-mer size of 20 was used for building the Meryl database from the raw CCS reads. Copy number k-mer plots were generated using Merqury’s provided R scripts. BUSCO version 5.1.2 (Manni et al. 2021) was also used in genome mode to assess genome ortholog completeness using the Lepidoptera OrthoDB-10 database (Kriventseva et al. 2019).

To detect contigs that were contaminant DNA and not of *T. absoluta* origin, blastn version 2.12.0 (Camacho et al. 2009) was used with the “nt” database (downloaded August 3^rd^, 2021) under the following parameters: word size=20, max target sequences = 10. Taxonomic information was downloaded for each subject match from the NCBI Taxonomy database using the “rentrez” package in R. To corroborate blast results, Phyloligo version 1.0 (Mallet et al. 2017) was used to generate a Euclidian distance matrix between contigs based on k-mer distribution with k-mer length 4. The R packages “ape” version 5.6-1 (Paradis & Schliep 2019) and “ggtree” version 2.2.4 (Yu et al. 2017) were used to generate and visualize the contig tree using the BIONJ algorithm.

### Genome Annotation

#### Repeat masking

The decontaminated hifiasm primary genome assembly was supplied as a database to RepeatModeler version 2.0.2a (Flynn et al. 2020) to produce a custom repeat library, with the long terminal repeat module enabled. RepeatMasker version 4.1.2 (Smit et al. 2021) was used to produce GFF annotation files of repeat coordinates. The Dfam transposable element database provided with RepeatMasker was merged with the RepBase RepeatMasker Edition database version 20181026 (Bao et al. 2015) to mask the genome once, and the custom RepeatModeler library was used to mask the genome separately. The resulting GFF files were merged and sorted with bedtools version 2.30 sort (Quinlan & Hall 2010), then used to soft-mask the assembly with bedtools maskfasta.

#### Gene model annotation

Six RNAseq datasets produced by Camargo *et al*. 2015 covering all life stages of *T. absoluta* (egg, four larval instars, adult) were downloaded from Bioproject PRJNA291932 for gene model annotation. Reads were checked for quality with FastQC version 0.11.9 (Babraham Bioinformatics 2019), trimmed with Trimmomatic version 0.39 (Bolger et al. 2014), and aligned to our primary assembly using STAR version 2.7.9a (Dobin et al. 2013) with default parameters. In addition, protein databases were downloaded from Lepbase (Challi et al. 2016) and OrthoDB-arthropoda (Kriventseva et al. 2019). The soft-masked genome was annotated twice with BRAKER2 version 2.1.5 (Brůna et al. 2021), once with the aligned RNA data, and again with the merged protein data. The two resulting GTF gene model files, as well as the GFF gene model hints files, were supplied to TSEBRA version 1.0.2 (Gabriel et al. 2021) to be merged into a single GTF output. Gffread version 0.12.6 (Pertea & Pertea 2020) was used to remove mRNAs with missing start or stop codons, in-frame stop codons, or that were redundant.

#### Functional gene annotation

Entap version 0.10.7 (Hart et al. 2020) was used to annotate gene models with names and predicted functions. Entap was configured to perform frame selection and filtering. The Lepbase, Refseq-Invertebrate, UniprotKB/Swiss-Prot, and UniprotKB/TrEMBL (O’Leary et al. 2016; Challi et al. 2016; The UniProt Consortium 2021) protein databases were used for gene identity search and the EggNOG database (Huerta-Cepas et al. 2019) for gene ontology, protein domain, and pathway annotation. As EnTAP was designed for transcriptome annotation, gene model coordinates were not referenced to the genome assembly. To correct this, the output GFF gene model annotation was converted to a GTF using gffreads, then converted to an alignment-GFF format using the script “gtf_to_alignment_gff3.pl” from Transdecoder (Brian & Papanicolaou), and finally mapped back to the genome assembly coordinates using the Transdecoder script “cdna_alignment_orf_to_genome_orf.pl”.

### Population sample DNA extraction and alignment

We used the same DNA extracted from *T. absoluta* collected from South America, Costa Rica, and Spain by Tabuloc *et al*. 2019. DNA libraries were made using the KAPA Hyperplus Kit (Roche, Basel, Switzerland). 150 basepair paired-end sequencing was performed by Novogene on the Illumina HiSeq 4000. Raw reads were trimmed of adapter sequences using scythe version 0.991 (Buffalo 2014) and were quality-filtered using sickle version 1.33 (Joshi & Fass 2011) using default settings. FastQC was used to inspect read quality before and after filtering. Reads were mapped to the genome assembly using bwa mem version 0.7.17 (Li 2013), and duplicates were marked with samtools markdup version 1.14 (Danecek et al. 2021).

### Population structure analysis

Angsd version 0.935 (Korneliussen et al. 2014) was used to estimate genotype likelihoods from the 78 contigs longer than 100kb, using a single nucleotide polymorphism (SNP) filter threshold of p < 10^−6^, a minimum minor allele frequency of 0.05, and a minimum map and base quality of Q=20. SNPs were then pruned to every 500 base pairs. PCA and admixture analysis were performed using PCAngsd version 1.0 (Meisner & Albrechtsen 2018) and NGSadmix version 32 (Skotte et al. 2013) as described in Lewald *et al*. 2021. PCAngsd was also used to output inbreeding coefficients for each sample.

### Treemix

We called genotypes from the genotype likelihoods calculated for population structure analysis using PCAngsd, with a 95% confidence threshold and with inbreeding values estimated from PCAngsd as priors. We removed loci that were missing data in more than 20% of samples, or loci with missing data in all individuals within a single sampling location. A custom R script was used to convert PCAngsd-format genotypes into a Treemix-formatted allele counts table. Treemix version 1.13 (Pickrell & Pritchard 2012) was run with 100 bootstraps, a window block size of 500 SNPs, 0 to 5 migration edges, and rooted on the CR (Costa Rica) population. The “global rearrangements” and “standard error calculation” options were also enabled. Treemix’s “threepop” subprogram was used to calculate F3 statistics between populations, using a 500 SNP window block size for standard error estimation.

### Population summary statistics and Population Branch Statistic

Summary statistics were calculated with Angsd on the largest 78 contigs. The site allele frequencies (SAF) were calculated for each region (North, Andes, Central, and Spain) with the -doMaf 1 option and using individual inbreeding coefficients as priors, and no minor allele frequency or SNP filtering was used. To estimate nucleotide diversity and Tajima’s D, the global folded 1-dimensional site frequency spectrum (1D-SFS) was calculated using Angsd realSFS for each population using the SAF in 100Mb pieces of the genome with a maximum of 400 iterations in the EM cycle. The 1D-SFS was summed across the genome for each population and realSFS saf2theta was used to estimate thetas per site. Angsd thetaStat do_stat was then used to calculate theta and Tajima’s D in 20kb windows, with a step of either 20kb or 5kb.

To estimate Fst between regions, realSFS was used to calculate the global folded 2-dimensional site frequency spectrum (2D-SFS) between each pair of regions in 100Mb pieces of the genome with 400 EM cycle iterations. 2D-SFS was summed across the genome, and realSFS Fst index was then used to estimate per-site Fst, and realSFS fst stats was used to estimate the global Fst values between regions. To estimate the Population Branch Statistic (PBS), the Andes, Central, and North regions were supplied at once to realSFS Fst index and realSFS fst stats2, to produce PBS values for each region in sliding 5kb windows with a 500bp step across the genome. Windows with less than 4kb of sequence data were excluded from the analysis. To compare allele frequencies of key SNPs between populations, we repeated site allele frequency estimation, but forced the reference allele to be the “major” allele.

### Population Modeling

The 2D-SFS was estimated between North, Andes, and Central regions using the same procedure as for summary statistics but excluded all genic regions (gene model boundaries plus an additional 1kb on flanking sides). Additionally, the Villa Alegre, Chile (VA) population samples were excluded from the analysis. Output 2D-SFS was converted to the Fastsimcoal version 2708 (Excoffier et al. 2013) format using a custom R script.

Fastsimcoal was used to estimate model parameters from the 2D-SFS data under several possible models. In the simplest model, an ancestral population is allowed to split two times, each with its own population size (5 total parameters). The exponential growth model adds a growth event to each population with unique exponential growth rates (9 total parameters). Finally, the resized population model replaces the exponential growth with an instantaneous change in population size (11 total parameters). For each model, parameter estimation was run 100 independent times, using 1 million simulations and 100 EM loops per run. SFS categories with less than 10 counts were excluded.

To compare models to each other and the data, 1Mb of DNA was simulated 100 times under each model using Fastsimcoal with a mutation rate of 2.9×10^−9^ basepairs/generation (Keightley et al. 2015) and a recombination rate of 2.97×10^−8^ cM/Mb (Yamamoto et al. 2008), based on estimates from *Heliconius melpomene* and *Bombyx mori*, respectively. The resulting genotypes were used to estimate the r^2^ - measure of linkage disequilibrium (LD) between SNPs less than 10kb away using ngsLD version 1.1.1 (Fox et al. 2019). To estimate LD from the sequencing data, the same minor allele frequency files used for Fastsimcoal parameter estimation were used to call genotype likelihoods in each region on the largest 29 contigs. NgsLD was run on a 10% subset of these genotype likelihoods to a max distance of 10kb.

Estimates of LD decay rates, maximum, and minimum LD were calculated from a 1% subset of r^2^ values from simulations and a 10% subset of r^2^ values from data using the provided fit_LDdecay.R script with the following parameters: fit_bin_size = 100, recombination rate = 2.97, fit_boot = 100, fit_level = 10.

### Data availability

Scripts used for genome assembly and population analysis available at github.com/ClockLabX/Tabs-genome-popgen. Genome assembly available at BioSample ID SAMN32102173.

## Results

### New Pacbio *Tuta absoluta* genome assembly improves gene annotation and contiguity

Before performing any population analyses, we decided to produce a high-quality reference genome based on long-read technology. New protocols for PacBio HiFi sequencing allow for low DNA input, which was critical in our case as Lepidopterans are notoriously heterozygous and using DNA pooled from many individuals would make genome assembly challenging. We sequenced a single moth originating from a laboratory colony at the Institute of Agrifood Research and Technology (IRTA), Cabrils, Spain, and obtained 16.2Gbp of sequence after collapsing circular consensus reads. Based on k-mer analysis with GenomeScope, the genome is 2.9% heterozygous and has 38.8% repeat content. GC vs k-mer plots show that there is likely no mass contamination from other species or microbes (Figure S1). The k-mer based haploid length estimate is 524 Mbp, which is close to the 564 Mbp estimate based on flow cytometry (Paladino et al. 2016).

To assemble reads, we used the HiFi assembler hifiasm and compared quality and contiguity metrics using Merqury. The accurate long reads allow for the ability to separately assemble maternal and paternal haplotypes at heterozygous regions. While hifiasm attempts to separate assembled contigs into primary and alternate haplotypes, we found that the primary assembly still had high haplotype retention based on its 990 Mbp length and the large peak of raw read k-mers that appear twice in the assembly (Figure S2A). We decided to further remove haplotigs using purge_dups, which shrunk the primary assembly size to 650.6 Mbp and eliminated the 2x raw read k-mer peak (Figure S2B).

Additionally, BUSCO analysis using the OrthoDB Lepidoptera gene set found the percent of complete, duplicated BUSCOs dropped from 48.5% in the unpurged assembly to 6.2% in the purged assembly (Figure S2C). This means that improperly retained alternate haplotypes have been removed from the primary assembly. When we examine the raw read k-mer multiplicity in the primary assembly, we see a peak of k-mers that map only to the primary or alternate assembly at k=13, which corresponds to the heterozygous portions of the genome (Figure 1B). We also see a peak at k=26 in k-mers that are shared between the primary and alternate assemblies, which matches the expectation of a diploid genome with double the read coverage in any homozygous regions of the genome. The frequencies of k-mers missing from either assembly are low, and represent k-mers from sequencing read errors.

**Figure 1:**
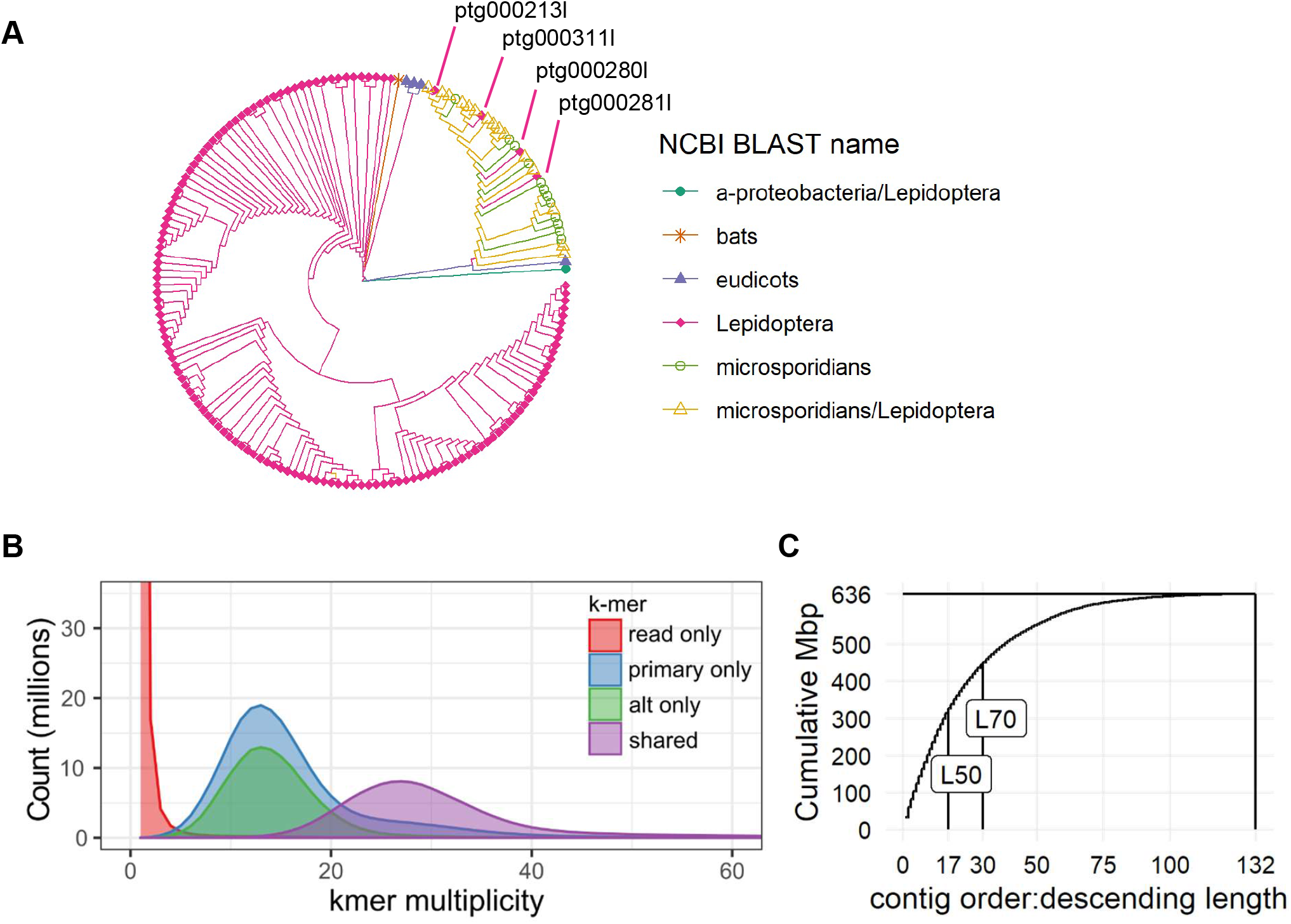
Genome assembly assessment. (A) K-mer based distance tree of contigs in the primary genome assembly before decontamination, labeled by NCBI BLAST matches in the “nt” database. Contig ptg000311l matched to insect mitochondrial sequences. Contigs ptg000311l, pg000280l, and ptg000281l were contigs annotated as “Lepidoptera” by BLAST but clustered with microsporidian sequences in the tree. (B) K-mer multiplicity plot of input CCS reads against the assembly. “Read only” indicates k-mers that only appear the raw reads; “primary only” “alt only” indicate read k-mers that appear in only one of the 2 haplotig sets; “shared” indicates k-mers that appear in both haplotig sets. (C) Cumulative length of contigs in primary assembly, ordered from longest to shortest. L50 and L70 indicate smallest number of contigs that contain 50% or 70% of the assembly length.

To identify which contigs came from contaminant DNA, we used the BLAST nt database, as well as a k-mer distance tree (Figure 1A) and found multiple contaminants. We identified a *Wolbachia* genome contig, several tomato contigs, and many microsporidian contigs, primarily from the *Nosema* genus (a common insect fungal parasite), all of which were expected. We also identified multiple contigs that matched to both *Nosema* and Lepidopteran queries in BLAST. Based on the distance tree’s clustering of these contigs with other *Nosema*-only contigs, as well as the GC content (Figure S3A), we decided to exclude these from the assembly. We also noticed four contigs that matched only to Lepidopteran contigs but clustered with Nosema sequences in k-mer content. One of these, ptg000311, matched to Lepidopteran mitochondrial sequences and likely represents the *T. absoluta* mitogenome. The remaining three matched to the same *Papilo xuthus* genome assembly (PRJNA291600) and is likely the result of inaccurate annotation in the BLAST database, as no decontamination steps were taken during its assembly (Nishikawa et al. 2015). In addition, these contigs’ GC and repeat content profiles were distinct from all other Lepidopteran contigs (Figure S3), so we excluded them from the assembly. Finally, one contig matched to a mouse-eared bat (*Myosis)* mitochondrial genome; this was possibly contamination from the sequencing facility and was excluded as well.

After removing these contigs, our primary assembly contained 132 contigs all longer than 10 kb with a final length of 635.9 Mbp. 70% of the genome was captured in the 30 longest contigs (L70); as *T. absoluta* has 29 chromosomes, this suggests our assembly is approaching chromosome-level contigs (Figure 1C). This represents a significant improvement from the previously published *T. absoluta* genome which consists of 81,653 contigs and a length of 906 Mbp (Tabuloc et al. 2019).

To generate gene models and functionally annotate genes, we used RepeatModeler and RepeatMasker to soft-mask the genome for repeats using both known Lepidopteran repeat sequences, as well as *de novo* sequences identified from our assembly. We followed with BRAKER2 to identify gene models, and functionally annotated models with EnTAP, based on multiple protein databases (RefSeq Invertebrate, UniProt SwissProt and TrEMBL, Lepbase, and EggNOG) and published *T. absoluta* RNAseq datasets. Of the 19,570 gene models identified by BRAKER2, 17,183 were identified as complete by gffread and EnTAP, with 14,019 transcript models matching to a gene name or functional annotation.

### Population Structure of *Tuta absoluta*

#### Population sampling and sequencing

We analyzed whole-genome sequencing data from individuals previously collected from field and greenhouse sites across South America and Costa Rica, as well as a lab colony from Spain (Tabuloc et al. 2019) (Figure 2A). Mapping rates to the new genome assembly ranged between 70%-90%, although sequencing depth per individual was low (between 1X to 17X) (Figure S4). One population from Argentina (MP, Mar del Plata, Buenos Aires State) had extremely low mapping rates and read depth, so we excluded it from further analysis. Wherever possible, we used methods based on genotype likelihoods, rather than genotype calls, to account for uncertainty that results from the low read depth.

**Figure 2:**
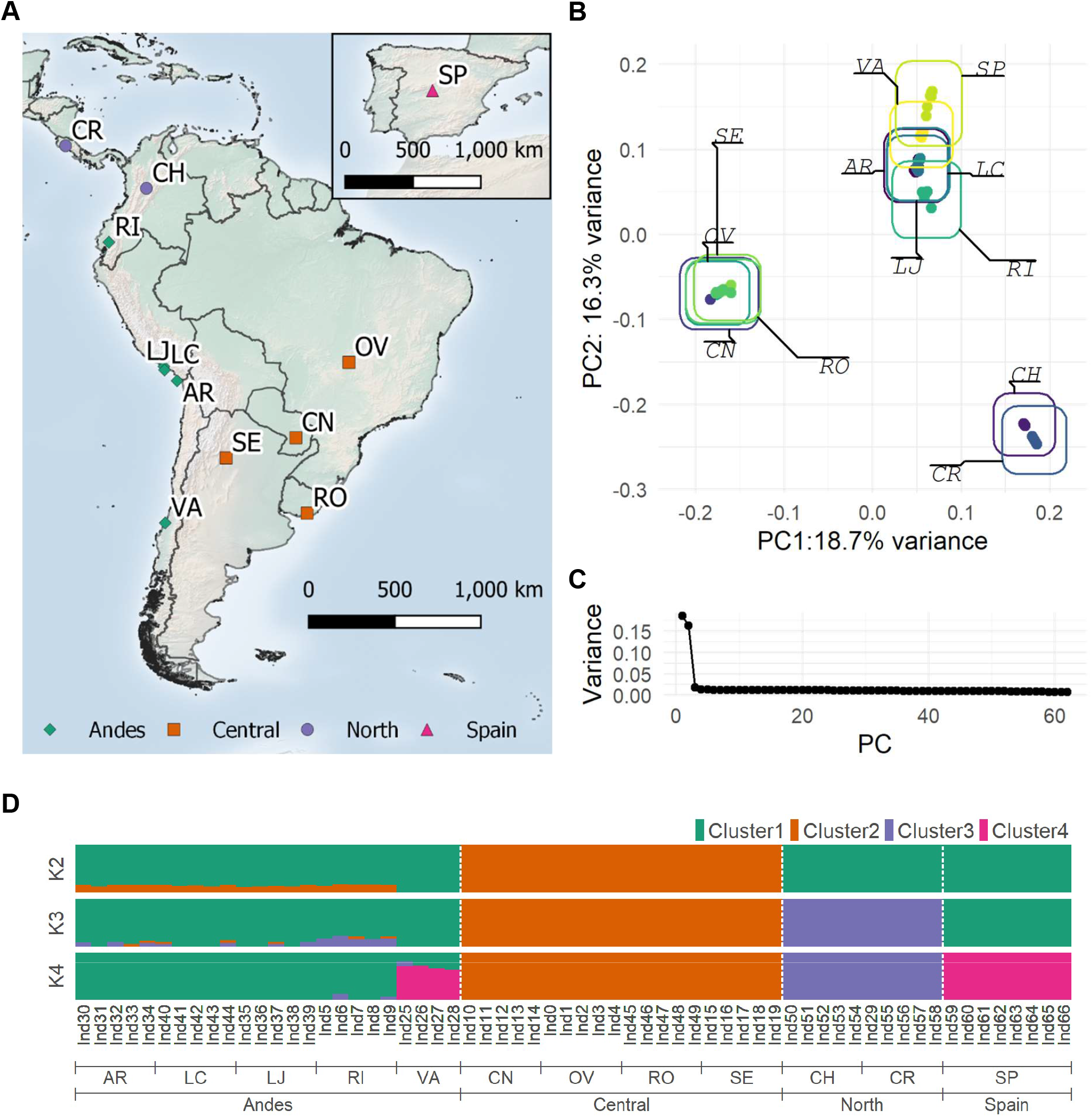
Sampling sites and population structure of T. absoluta individuals. (A) Map of sampling locations. Between 4 to 8 individuals were sequenced per location. Legend indicates grouping as determined from PCA and admixture analyses. (B) PCA plot of the first two principal components (PCs), based on genotype likelihoods from 933,060 sites. (C) Percent variance captured by PCs in descending order. (D) Admixture analysis for 2 to 4 clusters. Colored bars indicate the posterior probability of an individual belonging to a given cluster. Location codes: AR=Arica, Chile; CH=Chia, Columbia; CN=Campo Nove, Paraguay; CR=Costa Rica; LC=La Curva, Peru; LJ=La Joia, Peru; OV=Ouro Verde, Brazil; RI = Riobamba, Ecuador; RO=Rocha, Uruguay; SE = Santiago del Estero, Argentina; SP = Barcelona, Spain; VA = Villa Alegre, Chile.

#### Three distinct *Tuta absoluta* populations exist in Latin America

To investigate population structure in our samples, we used Principal Component Analysis (PCA) and admixture estimation based on allele frequencies from over 900,000 SNPs. The first two PCs captured 18.7% and 16.3% of the total variance in the data, with the remaining PCs each capturing less than 5% of the total data variance (Figure 2C). Samples primarily cluster together based on collection site but also formed three distinct regional groups (Figure 2B). Samples from Chile, Peru, and Ecuador form an “Andes” cluster west of the Andes Mountains; samples from Brazil, Uruguay, Paraguay, and Argentina form a “Central” cluster, east of the Andes Mountains; and samples from Columbia and Costa Rica form a “North” cluster. Spanish samples grouped tightly with the Andes cluster, particularly the VA (Villa Alegre, Chile) site.

When three clusters were allowed in admixture estimation, samples group into the same three clusters as in PCA, while at four clusters, the Spanish samples become their own group, with VA samples sharing a large proportion of admixture. Compared to other Andes populations, the VA samples are more differentiated from Central and North sites as well, with little signal of admixture at all levels of k tested. The other Andes populations (AR, LC, LJ, and RI) all had low admixture proportions from Central at k=2, although at k=3 we see that all RI (Riobamba, Ecuador) samples exhibited admixture from the North populations. This suggests that the non-VA Andes populations are more closely related to Central populations than VA, and that VA could represent an admixture between the population that gave rise to the Spanish lineage and the other Andes populations. Additionally, we see that RI represents an intermediate population between the Andes and North, which makes sense given its geographic location between the two clusters.

To further quantify population structure between these clusters, we calculated nucleotide diversity, Tajima’s D, and Fst using genotype likelihoods (Figure S5). For all clusters, nucleotide diversity was approximately 2%, which is fairly high compared to most Lepidopterans (Mackintosh et al. 2019). If we look at the weighted Fst, we see differentiation between clusters is high, particularly between North and all other clusters. The combination of high diversity levels and high Fst could mean these regions diverged from each other a long time ago, prior to the detection of *T. absoluta* by growers across Latin America in the 1960s to 1980s. If divergence had occurred recently, we might expect reduced diversity levels in invasive populations relative to the ancestral population.

#### Treemix confirms clustering and detects migration events to Ecuador and Chile

To detect potential migration events between populations, we used Treemix to build a maximum likelihood (ML) tree based on allele frequencies, as well as predict migration edges and calculate F3 statistics. As Treemix was designed to take allele count data per population, we called genotypes using PCAngsd using a 95% accuracy cutoff and counted alleles within each sampling location. After filtering out loci with missing data, 47,535 SNPs were available for use. In general, the tree topography aligns with results from PCA and admixture analyses. We see sampled sites cluster into the same three clusters, North, Andes, and Central, with the Spanish samples sister to the VA (Chile) site (Figure 3). In agreement with Fst estimates, North populations have experienced more genetic drift from the Andes and Central populations, compared to the Andes and Central populations with each other.

**Figure 3:**
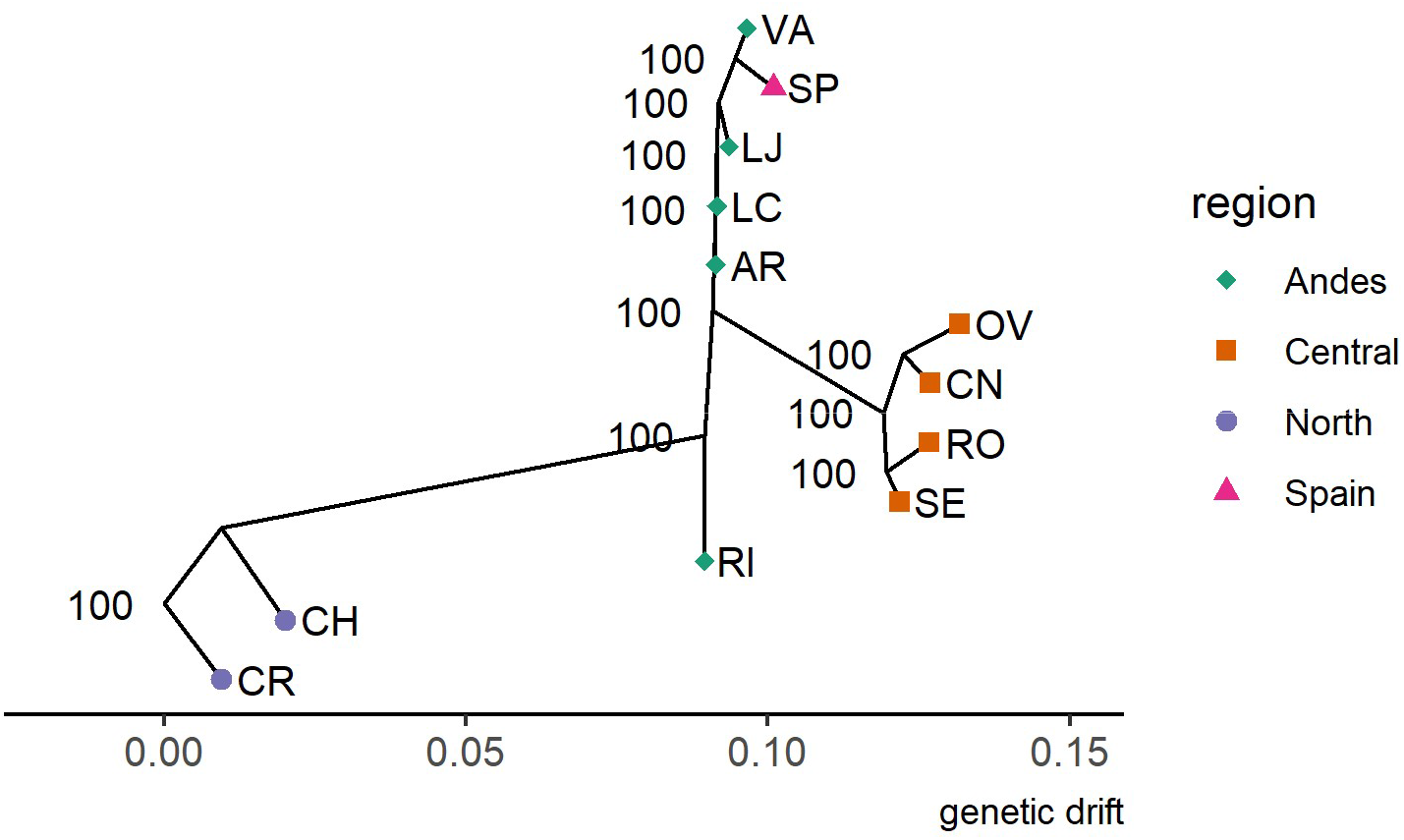
Maximum likelihood tree from Treemix with no migration edges called rooted on the Costa Rica (CR) samples, based on 47,535 SNPs. Confidence values were based on 100 jackknife bootstraps with 500 SNP bins. X-axis represents genetic drift distance between populations.

**Figure 4:**
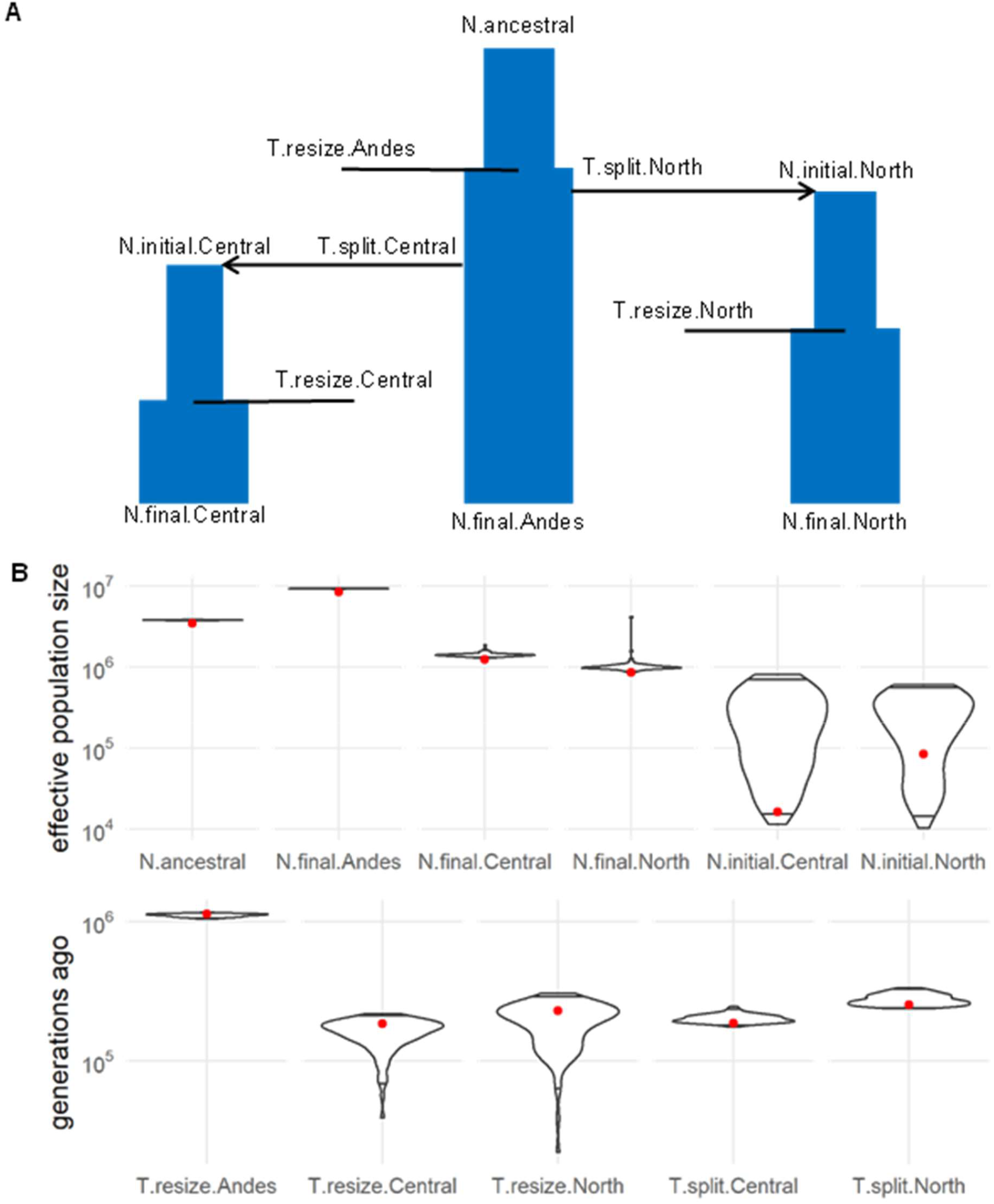
Coalescent simulations used to estimate model parameters. (A) Model for a three-population split with population resizing events for each population. (B) Maximum likelihood estimates of each parameter in the model indicated by a red dot. Violin plot depicts distribution of parameter values from 100 parametric bootstraps, with upper and lower boundary lines indicating the 95% interval. All point estimates were within the 95% bootstrap intervals except N.ancestral, N.final.Andes, N.final.Central, and N.final.North.

Interestingly, the RI (Ecuador) site does not form a clade with other Central populations but descends from the common ancestor of the Central/Andes group. Based on admixture analysis that showed low levels of admixture in RI from the North, the position of RI in the tree could be further evidence that Ecuador represents an intermediate mixing zone between populations north and south of it. To investigate further, we re-ran Treemix allowing between one to five migration events (m=0 to 5) and calculated the F3 statistic between all combinations of three populations to see if admixture was supported. At m=2,4 and 5, Treemix reported a strong migration (between 11%-25%) from the Spain or Spain/VA branch to RI, while at m=3 and 5, Treemix reported a weak migration event (2%-7%) from the North to RI. F3 statistics F_3_(RI; CH,SP) and F_3_(RI; CR,SP) were significantly negative (Table 2), indicating that a simple bifurcating tree does not explain RI’s relationship with CH, CR, and SP. While Treemix infers a migration from the Spanish branch to the RI branch, it is important to remember that this migration is inferred to have occurred somewhere along the branch between the current day Spanish population and the most recent common ancestor of Spain and VA (Chile). This migration could have occurred early in the branch, when the population was still in Chile, or late in the branch, when the population moved to Spain. Based on fresh tomato trade between the two countries, in 2006 Chile shipped over 29,000 kg of fresh tomatoes to Spain while importing none back (2006). This makes admixture from Spain back to RI unlikely and suggests that RI contains admixture from North and Chilean populations.

**Table 1:** Migration events with predicted weights from Treemix between populations, under different models allowing between 1 to 5 migration events (m). * indicates the migration significantly improved model fit (p < 0.05) based on the Wald statistic using jackknife estimates. Inferred migrations originate somewhere between the “start-node” and the start-node’s most previous branchpoint, and terminate somewhere between the end-node and the end-node’s most previous branchpoint. As an example, a migration starting from “SP” occurs somewhere on the branch between “SP” and the most recent common ancestor of “SP” and “VA”.

| m | Weight (%) | start-node | end-node |
| --- | --- | --- | --- |
| 1 | 8.8* | SP | AR |
| 2 | 11.2* | SP | AR |
| 2 | 15.1* | VA,SP | RI |
| 3 | 17.7* | SP | AR |
| 3 | 17.3* | SP | LC |
| 3 | 2.0* | CR,CH | RI |
| 4 | 10.6* | SP | AR |
| 4 | 25.4* | VA,SP | RI |
| 4 | 2.2* | OV | RI |
| 4 | 13.6 | SP | LJ |
| 5 | 21.3* | SP | LC,AR |
| 5 | 11.0* | SP | RI |
| 5 | 24.2* | SP | LJ |
| 5 | 6.7* | CR | RI |
| 5 | 2.9* | LC,AR | OV |

**Table 2:** Significant F3 statistics. Population comparisons for which F3 statistics were significant (Z score <-3) are reported.

| Pop(A;B,C) | F3 | SE | Z score |
| --- | --- | --- | --- |
| AR;CH,SP | -0.00165 | 0.000171 | -9.63768 |
| AR;CR,SP | -0.00178 | 0.000171 | -10.4413 |
| AR;CN,SP | -0.00078 | 0.00015 | -5.15567 |
| AR;OV,SP | -0.00082 | 0.000145 | -5.63104 |
| AR;RO,SP | -0.00076 | 0.000151 | -5.05319 |
| AR;SE,SP | -0.00077 | 0.00014 | -5.47246 |
| LC;CH,SP | -0.00146 | 0.000193 | -7.56011 |
| LC;CR,SP | -0.00154 | 0.000211 | -7.31075 |
| LC;OV,SP | -0.00054 | 0.00018 | -3.00356 |
| RI;CH,SP | -0.00285 | 0.000948 | -3.00128 |
| RI;CR,SP | -0.00296 | 0.000965 | -3.06668 |

In addition to admixture in RI, Treemix and F3 statistics also detected admixture in AR (Chile) from the Spanish population. At m=1,2,3, and 4, Treemix detected a migration edge from the Spanish branch to AR with migration weight varying between 9% to 17%. F3 statistics of AR and Spain with any population from North or Central resulted in a significantly negative value, providing strong evidence of a migration event from a Spanish ancestor like the signal detected with RI. This suggests that the admixture signal we see in AR is from the same Chilean population that gave rise to the Spanish invasion and RI admixture.

### Divergence of *Tuta absoluta* predates modern tomato agriculture

Based on the high levels of nucleotide diversity and Fst between the Andes, Central, and North clusters, we hypothesized that the three regions may have diverged many generations ago, before the appearance and detection of *T. absoluta* in agricultural crops throughout South America in the mid-20^th^ century. This would suggest a model in which *T absoluta* may have adapted from local, wild host plants to nearby tomato fields independently, rather than a single population that became adapted to tomatoes and was spread through human activity. To investigate this, we calculated the folded two-dimension site frequency spectrum (2D-SFS) between populations and estimated parameter values under various population models using maximum likelihood coalescent methods (Figure S6, Table S2). We excluded the VA samples from the Andes cluster to avoid potential modeling issues due to VA appearing to originate from a distinct ancestor than other Andes populations. The simplest model allows for two population splits with constant population sizes, while the exponential growth model adds an exponential growth rate to each population. As exponential growth may not be appropriate if divergence times are long, we also tested a model with a simple resizing event for each population at some point in time. We used a post-hoc comparison of simulated linkage disequilibrium decay rates between models to test model fit. We found that while all three models simulated decay rates within the 95% confidence interval of the Andes population data, none simulated decay rates that overlapped with Central and North decay rate estimates, although the resizing population model was closest. (Figure S7). The lack of fit suggests there are additional complex historical events not well captured in these models. Under the resizing population model, divergence of the North occurred 252,383 generations ago (95% CI: 243,535-326,583), followed by a Central population divergence 187,034 generations ago (95% CI: 181,668-235,424). Reports of *T. absoluta* generation times can be as high as 6 to 12 or more generations per year (2005; Mansour et al. 2018; de Campos et al. 2021), dating these divergence events to tens of thousands of years ago. This suggests that *T. absoluta* was already present across Latin America prior to the 1960s, and as tomato agriculture surged, adapted locally to the new host plant.

### Population Branch Statistic Screening identifies several peaks under selection

The Population Branch Statistic (PBS) is an Fst-based statistic that uses Fst data between 3 populations to calculate the population-specific allele frequency changes. Regions of the genome with abnormally high PBS may be under strong selective forces, causing the loci allele frequencies to change faster than expected by drift. We calculated PBS across the genome for all three populations and found several peaks in contigs 2, 9, 15, and 22 that were exceptionally high and broad, particularly in the North cluster (Figure 5A). The peak in contig 9 contained the gene *paralytic* (*para, T. absoluta* gene g15590), a neuronal sodium channel protein that is the active target of pyrethroid insecticides (Dong et al. 2014). While PBS peaks in the North population between 13.1 and 13.2Mb on contig009, we note that the allelic diversity was low in the Andes and Central clusters relative to the North (Figure 5B). We calculated allele frequencies of known resistance-inducing mutations in each cluster (Dong et al. 2014), and found one mutation, an alanine to leucine substitution at position 1014, was fixed in the Central and Andes, while at 41% frequency in the North (Figure 5C). In addition, we found low to intermediate frequencies of other resistance alleles, including M918T, T929I, V1016G, L925M, and I254T.

**Figure 5:**
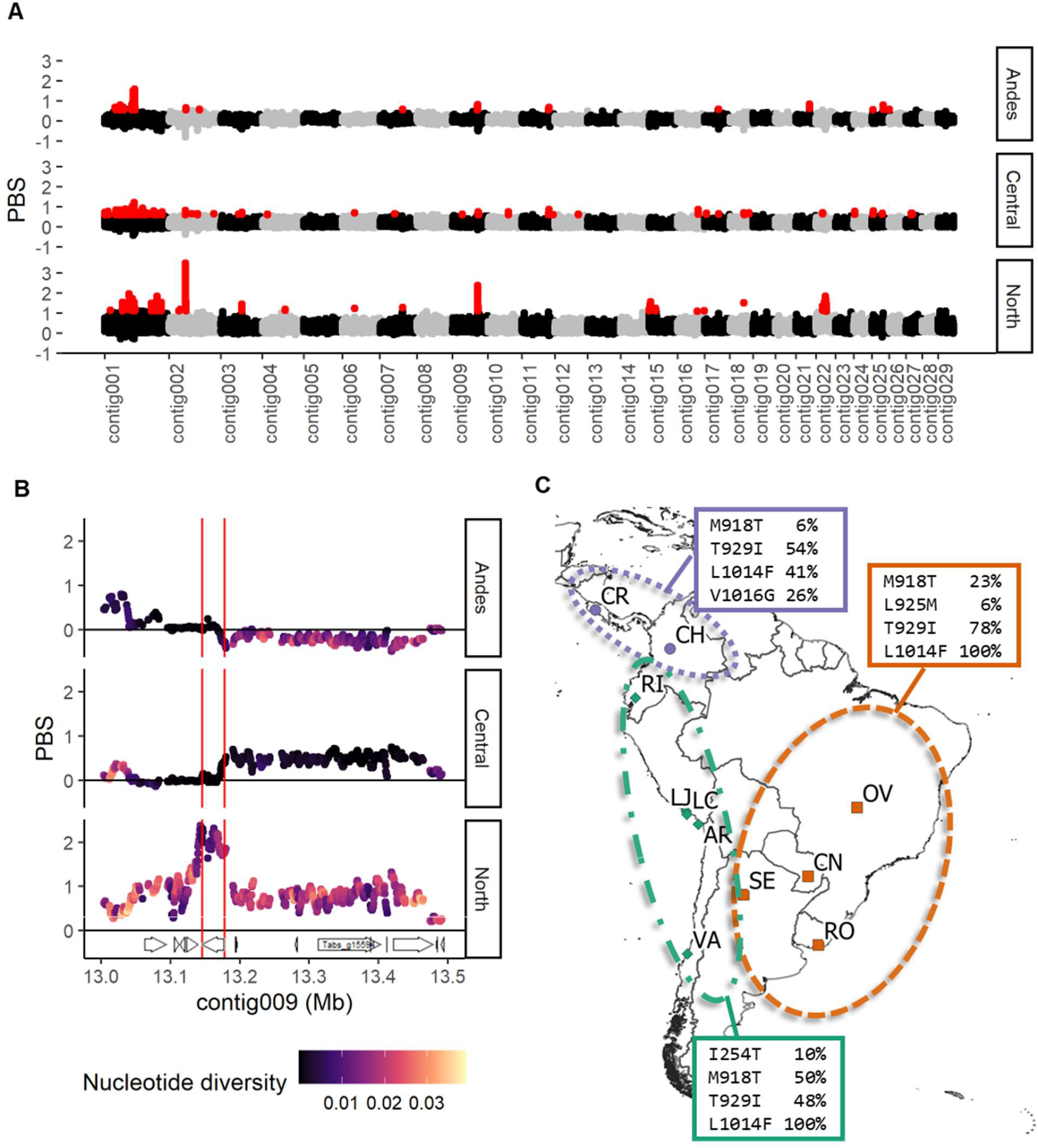
Selection signals in T. absoluta. (A) PBS values in each region calculated across the largest 29 contigs in 5kb intervals with 500bp steps. Red points indicate the highest 0.1% PBS values. (B) Plot of PBS and genetic diversity (pi) at contig009. Red vertical lines border the PARA gene model. (C) Map of sampling locations with allele frequencies of known amino acid substitutions that confer pyrethroid resistance within each cluster. Note the L1014F allele, which was found in all Central and Andes samples, while at 41% in North samples.

A similar selective sweep signal was also seen in the PBS hotspot on contig 2 (8.54Mb-8.58Mb), with high PBS and diversity levels in the North, and a large region of low allelic diversity in the Andes and Central (Figure S8A). Several genes were captured in this interval, including *NADH:Ubiquinone oxidoreductase subunit A8 (ndufa8)*; *mucin-5AC-like*, a gene putatively related to human *mucin-5AC*, and two hemomucin genes involved in hemocyte adhesion and innate immunity in insects. The PBS hotspot on contig 15 contained *cryptochrome-2 (cry2)*, encoding a key component of the circadian clock (Figure S8B). Finally, the large hotspot on contig 22 contained 69 genes, including multiple copies of *juvenile hormone binding protein*, ribosomal proteins, gustatory response genes, rhodopsins, and *acetylcholinesterase* (*ache*), a gene implicated in organophosphate insecticide resistance (Figure S8C).

Interestingly, based on alignment data it appears the Central and Andes populations may have two copies of *ache*, while North populations only have one. We looked for known mutations conferring organophosphate resistance and found moderate frequencies in all three clusters as well (Table S3).

## Discussion

Using whole genome sequencing data, we found that *T. absoluta* samples collected from 11 locations in Latin America clustered into three basic regions, comprised of a North, Andes, and Central group. In addition, we see that Spanish populations likely originated from a Central Chilean source based on their low level of Fst with the Andes and location on the ML tree. Previous analyses with mitochondrial sequences were unable to differentiate populations (Cifuentes et al. 2011); however, analyses using microsatellite data was able to identify these same three clusters and suggest a Central Chilean source for the European migration as well (Guillemaud et al. 2015). In agreement with this conclusion, looking at fresh tomato export data we see Chile is a worldwide exporter, shipping 12 tons of tomatoes an average distance of 11,700 km in 2018, while most other countries in South America tend to export within the continent (2018).

The Andes Mountains represent an obvious geographic barrier that would separate the Central population from the North and Andes populations. Population structure between Andes and North populations may be due to factors related to the changing latitude, including temperature and daylength. While PCA groups our Ecuador samples (RI) with Andes populations, admixture analysis and Treemix both provided some evidence that Ecuador may represent an admixture zone between the two regions. The weighted Fst between the North and Andes is also lower than between North and Central, suggesting that North and Andes are indeed more closely related. This general Fst pattern was also observed based on microsatellite analyses (Guillemaud et al. 2015). Sequencing of more samples from Peru and Ecuador might be needed to further elucidate the extent of an admixture zone between these clusters.

While *T. absoluta* was first discovered in Peru in 1917, its native range is not well established. One hypothesis is that *T. absoluta* migrated out of the Andes region and across South America through the 1960s-80s because of human transport by agricultural shipping. This aligns with the surge in domestic tomato agriculture in South America at the same time (Minami 1980). However, based on the similar nucleotide diversity levels between clusters, as well as high levels of Fst, we hypothesized it might be more likely that this migration across South America may have happened prior to tomato commercialization, with populations of *T. absoluta* later adapting to the appearance of commercial tomato agriculture. Based on our simple 3 population model, it appears the ancestral population diverged twice tens of thousands of years ago. Relative to the ancestral population size, the combined effective population size is roughly three times larger, although this is heavily weighted toward a very large Andes population, relative to the North and Central regions.

The fact that the estimated Andes population size is nearly 10 times larger than that of Central or North populations, as well as its slightly higher level of genetic diversity, could suggest that the Andes cluster represents the ancestral population range. Given that the wild ancestors of tomatoes and potatoes are also native to the Andes region (Spooner et al. 2005; Peralta & Spooner 2006), the Andes region would be an ideal place to search for native parasitoids for biocontrol. To date, biocontrol methods in South America have relied on non-natives or generalist parasites, with only a handful of native specialists identified in the literature and none that are commercially available (Salas Gervassio et al. 2019; Desneux et al. 2022). Thus, knowledge of *T. absoluta*’s native range may help focus efforts to identify more natural parasitoids.

Using the PBS, we found multiple genomic windows under apparent selective forces. Not surprisingly, one of the highest PBS windows contained the *para* gene, which encodes a sodium ion channel that is targeted by pyrethroids. The extremely low allele diversity in the Andes and Central populations relative to the North suggest a hard selective sweep occurred here. Heavy pyrethroid use in Brazil led to the appearance of resistant strains starting in the 1990s (Siqueira et al. 2000). We found that Central and Andes populations were completely fixed for the L1014F mutation, one of the most common causes of knock-down-resistance (kdr) to pyrethroids (Dong et al. 2014), while the North had an intermediate frequency. A study looking at Brazilian populations found a similar pattern of fixed L1014 (Silva et al. 2015), while another study looking at multiple populations in South America also found the same pattern of L1014F fixation in Central and Andes populations but not in the North (Haddi et al. 2012). While both studies also found M918T and T929I at elevated frequencies in all populations, we additionally detected the resistance allele V1016G in the North. We also found L925M in Central and I254T in Andes. While these have not been characterized as resistance alleles, mutations at these same positions have been shown to confer resistance in *Drosophila melanogaster* (I254N) and *Bemisia tabaci* (L925I) (Pittendrigh et al. 1997; Morin et al. 2002). These new appearances indicate selection is ongoing in all three clusters.

Other regions under selection were less obvious. We found one region in contig 2 containing *Ndufa8* and several hemomucin/mucin genes with a similar low genetic diversity in the Central and Andes populations and high PBS in North, indicating a hard selective sweep. *Ndufa8* produces a nuclear-encoded subunit of the NADH dehydrogenase complex I, part of the electron transport chain in the mitochondria used to generate ATP. Mutations here are known to cause mitochondrial complex I deficiency in humans (Yatsuka et al. 2020), although a few studies have found evidence of positive selection occurring in other species, potentially related to metabolism (Kozell et al. 2020; Lee et al. 2020). The hemomucin genes are a component of the insect immune system that are involved in endocytosis (Schmidt *et al*. 2010). Selection here could be in response to the increased use of parasitoids and predators such as *Trichogramma evanescens and Nesidiocoris tenuis* as an biological control alternative to insecticides (Han et al. 2019).

The elevated PBS region in contig 22 was relatively large at approximately 1Mb in size, containing over 60 genes. Interestingly, we noticed two copies of *ache* contained within this window, which codes for *acetylcholinesterase*, a gene which encodes a protein which degrades the neurotransmitter acetylcholine (Mutero et al. 1994). As this enzyme is the main target of organophosphates and carbamate insecticides, alleles in *ache* have been documented to confer resistance. Using the amino acid numbering scheme based on *T. california* (Massoulié et al. 1992), the resistance allele A201S has previously been reported to be present in European populations (Haddi et al. 2017), and we found this allele to be at moderate to high frequency in all three regions. We also found moderate frequencies of the mutation F290V and F290N. The F290V resistance allele has been documented in mosquitoes and moths (Cassanelli et al. 2006; Alout et al. 2009; Carvalho et al. 2013), while other mutations such as F290Y have been documented in *Drosophila* and *M. domestica* (Mutero et al. 1994; Walsh et al. 2001). Interestingly, based on mapping read depth the North population only contained a single copy of *ache*. As duplication of *ache* has been implicated in improved organophosphate resistance (Alout et al. 2009; Sonoda et al. 2014), this large structural duplication may be the reason for elevated PBS levels across such a large interval. Follow-up work with long-read methods or higher sequencing coverage will be needed to confirm the presence of structural duplication at this locus.

We expect that addition of a new contiguous genome assembly with annotations will be of benefit to the *T. absoluta* and Lepidopteran research community. Previous studies have worked to develop potential RNA interference (RNAi) strategies to use as an alternative to traditional pesticides (Camargo et al. 2015). Work is also being conducted to develop Cas9 gene-editing techniques for *T. absoluta* to facilitate future genetics studies (Ji et al. 2022). These developments in combination with an accurate assembly and gene annotations will allow for accelerated research towards understanding *T. absoluta* biology and methods to contain its economic impacts and spread.

## Contributions and Acknowledgements

K.M.L. and J.C.C. conceived the study. K.M.L. and C.A.T. performed nucleic acid extraction and library preparation for population samples. K.E.G., J.A., C.R.P., and J.C.G. provided field and colony collections. K.M.L. performed genome assembly and population analyses. K.M.L. and J.C.C. wrote the manuscript with input from all authors.

Thank you to Robert Munch (QB3, UC Berkeley) and John Alterio (PacBio, Menlo Park) for their advice and work on DNA extraction, library preparation, and long-read genome sequencing for the genome assembly. Thank you to Dr. Graham Coop and Dr. Jeffrey Ross-Ibarra for providing advice on population analyses.

This work is funded by NIFA 2020-67013-30976 and NIFA CA-D-ENM-2150-H.

## Supplemental Figures

**Figure S1:**
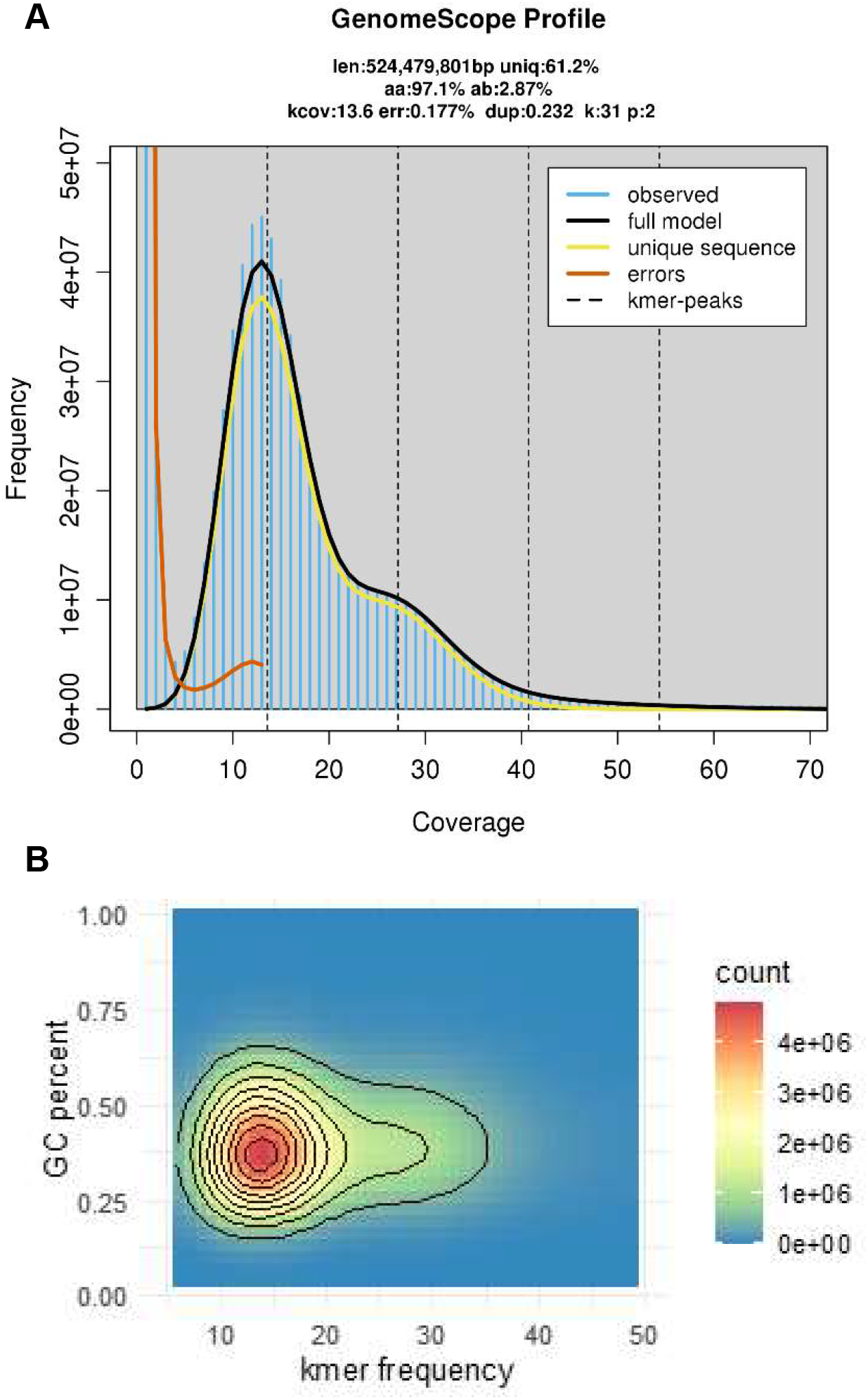
(A) GenomeScope profile of Pacbio CCS reads. (B) GC percent vs k-mer frequency plot of CCS reads, excluding k-mers with frequency less than or equal to 5 (to ignore unique k-mers due to sequencing errors).

**Figure S2:**
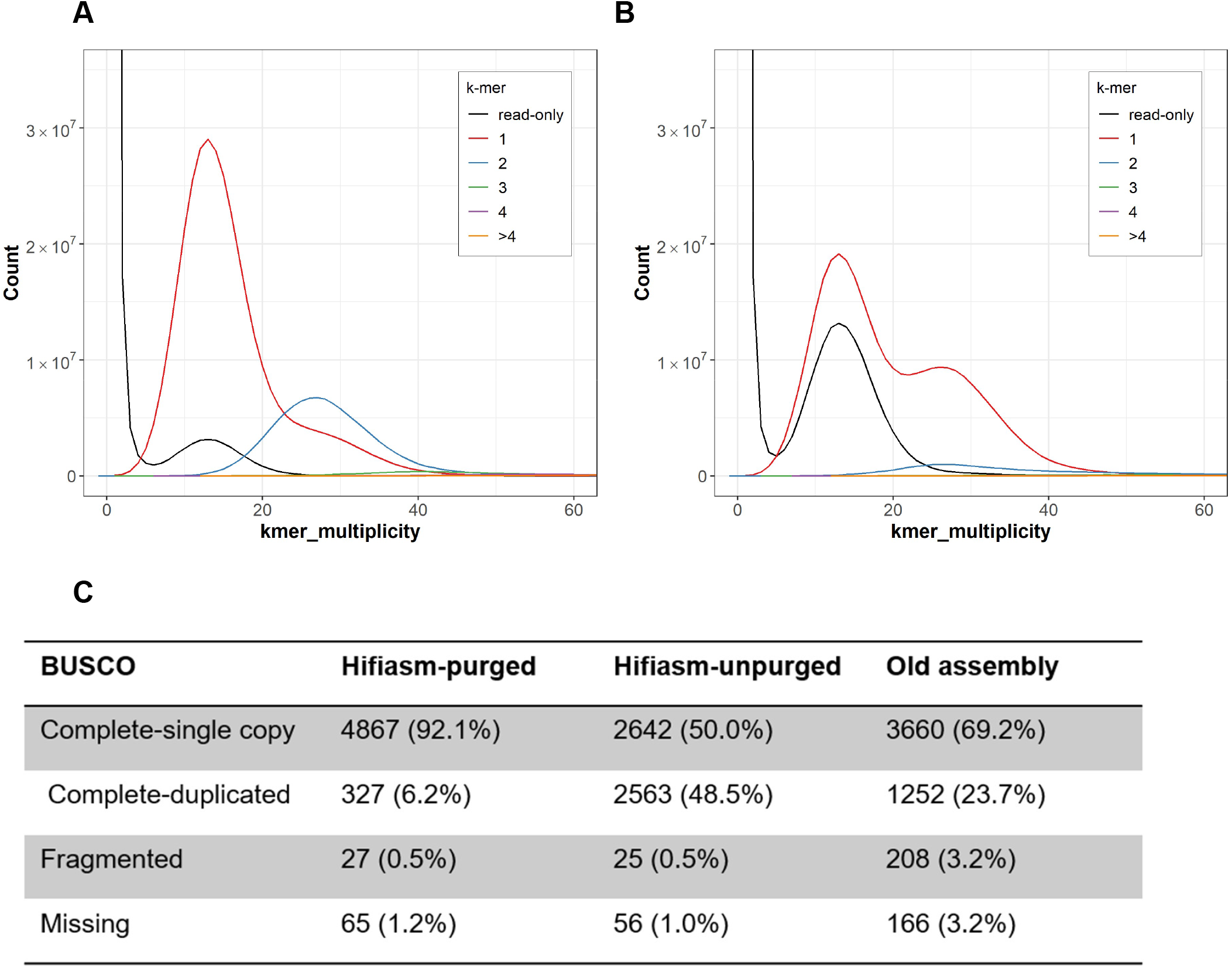
K-mer multiplicity plots by Merqury of the hifiasm primary assembly (A) before and (B) after purging retained haplotigs. (C) Table of Lepidopteran BUSCO scores of the purged and unpurged hifiasm primary assembly, compared to the previously published T. absoluta assembly (Tabuloc et al. 2019).

**Figure S3:**
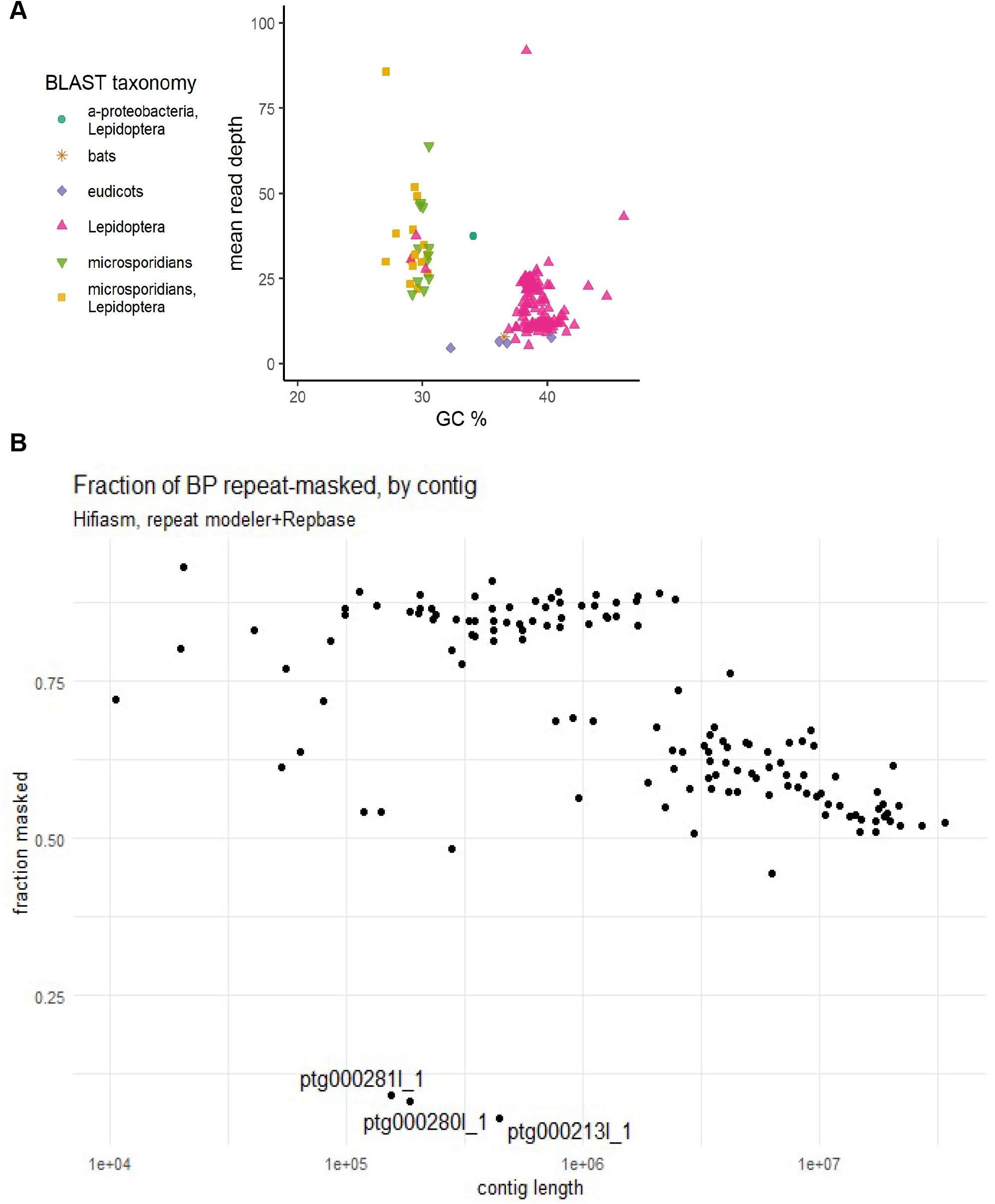
Decontamination results of the primary assembly. (A) Blobplot of primary contigs showing best BLAST matches vs GC % and mean read depth. The three “Lepidopteran” contigs at the 30% GC position were contigs ptg000311l, pg000280l, and ptg000281l, which appear to be microsporidian contamination that has been mislabeled as Lepidopteran. B) Repeat content of each contig in the primary assembly. Note contigs ptg000213, ptg000280, and ptg000281 have abnormally low repeat content.

**Figure S4:**
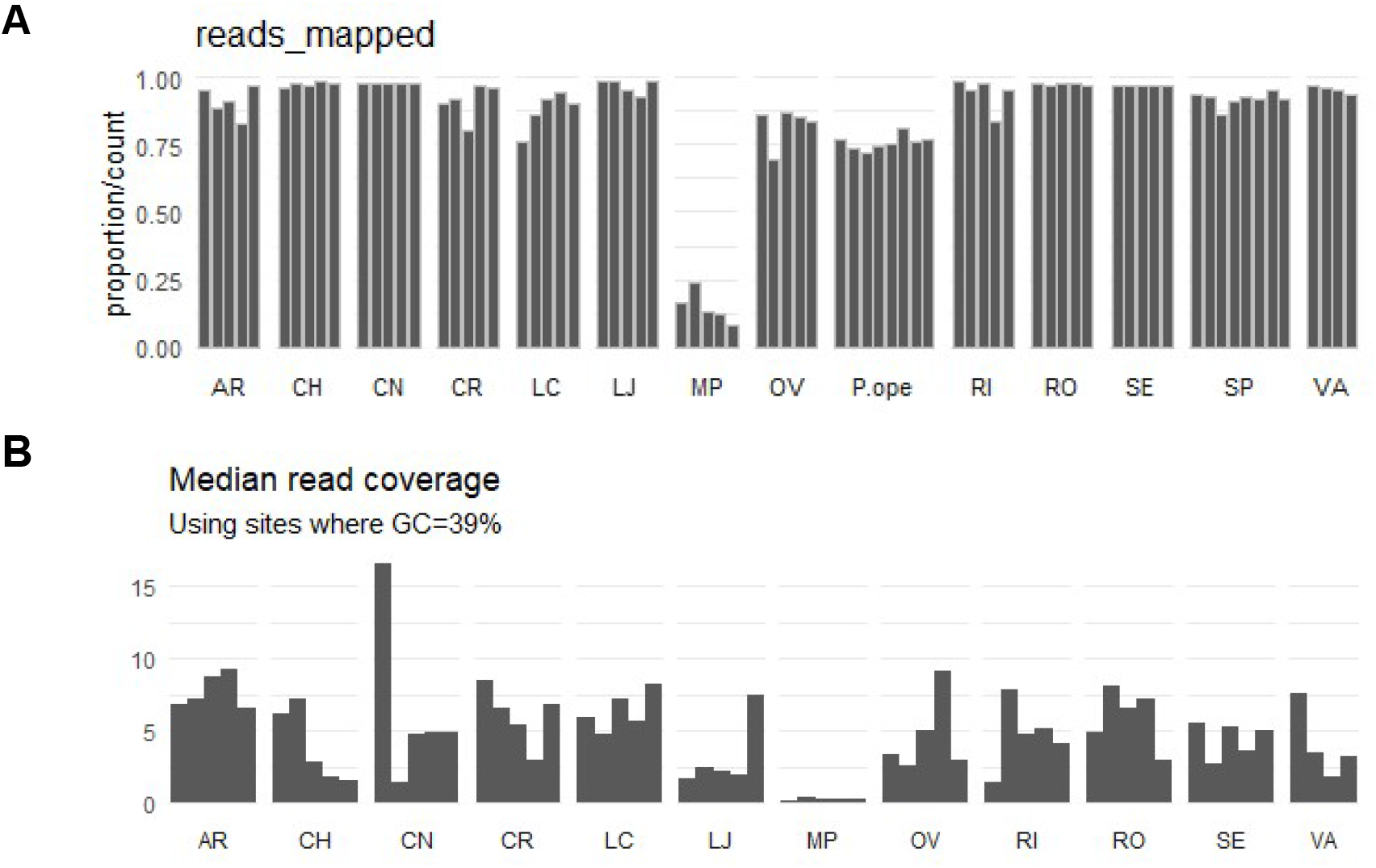
(A) Mapping rates and (B) median read coverage for Illumina reads from population sampling. Median read coverage was calculated at GC=39%, as this is the average GC content of the genome.

**Figure S5:**
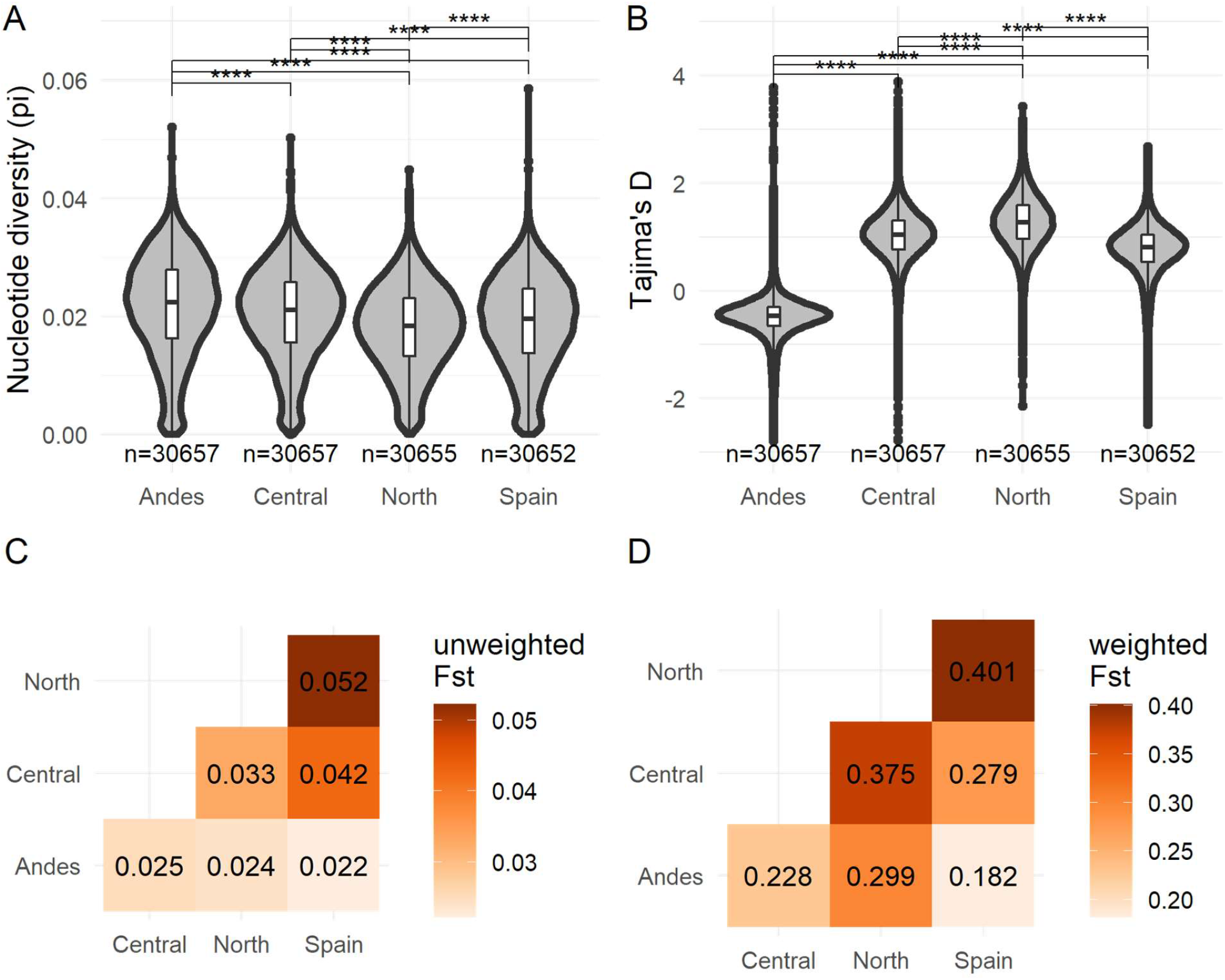
Summary statistics for the 4 key populations. (A) Pairwise nucleotide diversity and (B) Tajima’s D for each cluster, averaged across the genome in 20kb windows. N represents the number of windows included. Pairwise t-tests with a Holm’s correction method were used to compare means. **** indicates p-value < 0.0001. (C) Unweighted Fst and (D) weighted Fst calculated by Angsd. Weighted Fst is typically considered more accurate as it is less biased when using many rare, population-specific SNPSs (as is the case when genotyping by whole-genome sequencing) (Bhatia et al. 2013).

**Figure S6:**
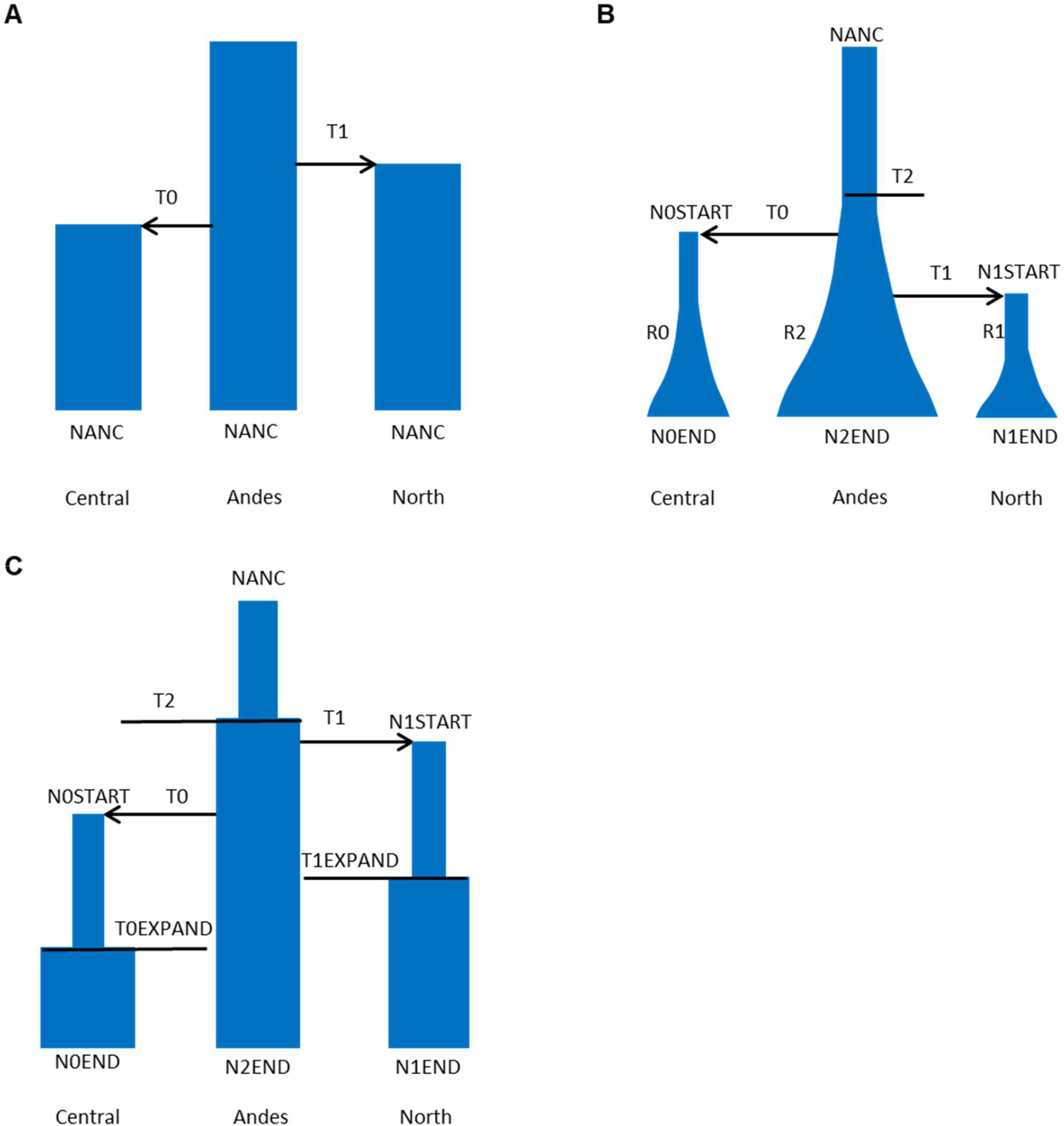
Three population models used to estimate parameter values. (A) M1: Model with two population splits and constant population size. (B) M2: Model with exponential growth. (C) M3: Model with population resizing events instead of exponential growth.

**Figure S7:**
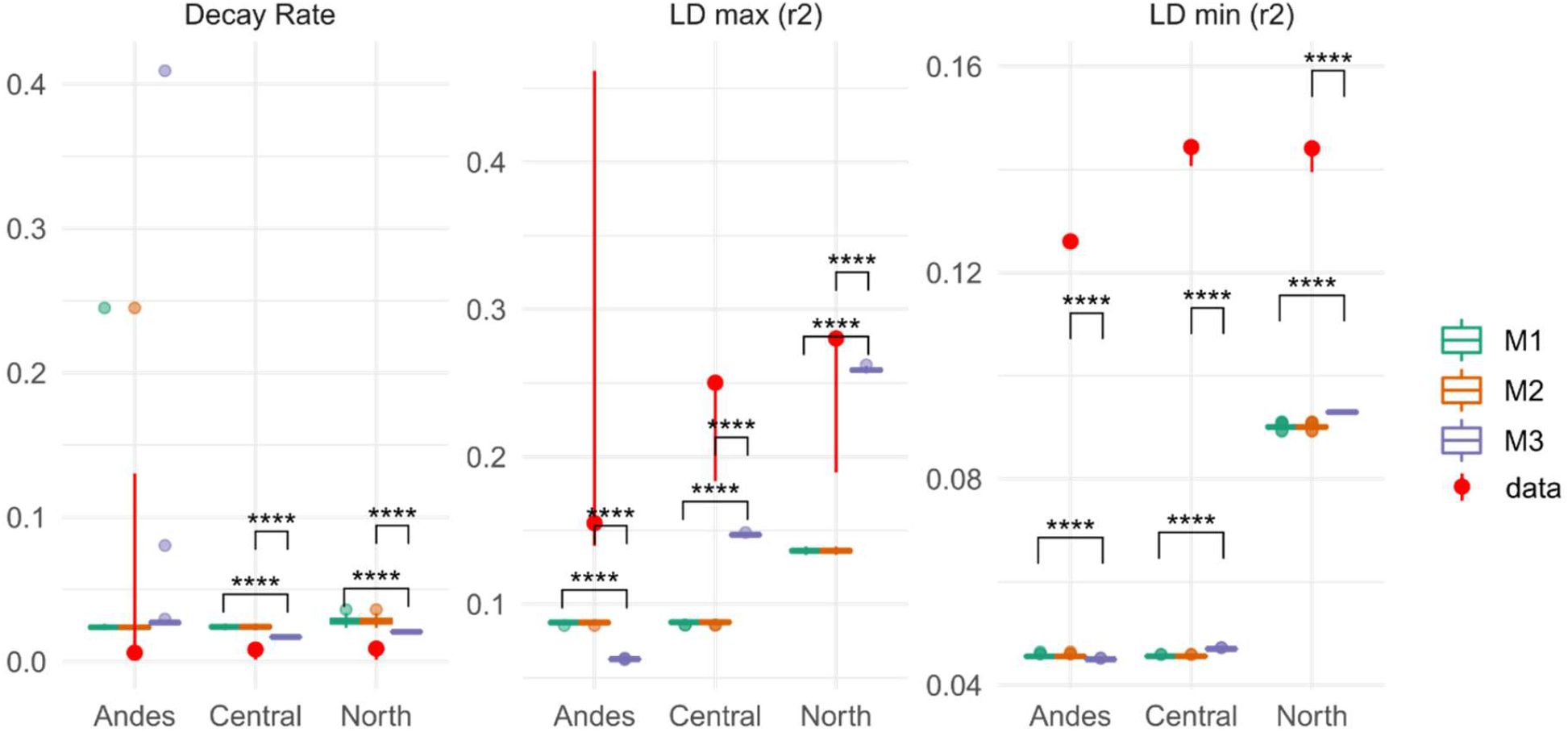
Linkage disequilibrium decay rates, minimums, and maximums over a 10kb interval calculated from genotype likelihoods or from 100 independent simulations under three different population history models. Values estimated from the data are shown by the red points with 95% confidence intervals. Pairwise t-tests with a Holm’s correction method were used to compare means. **** indicates p-value < 0.0001.

**Figure S8:**
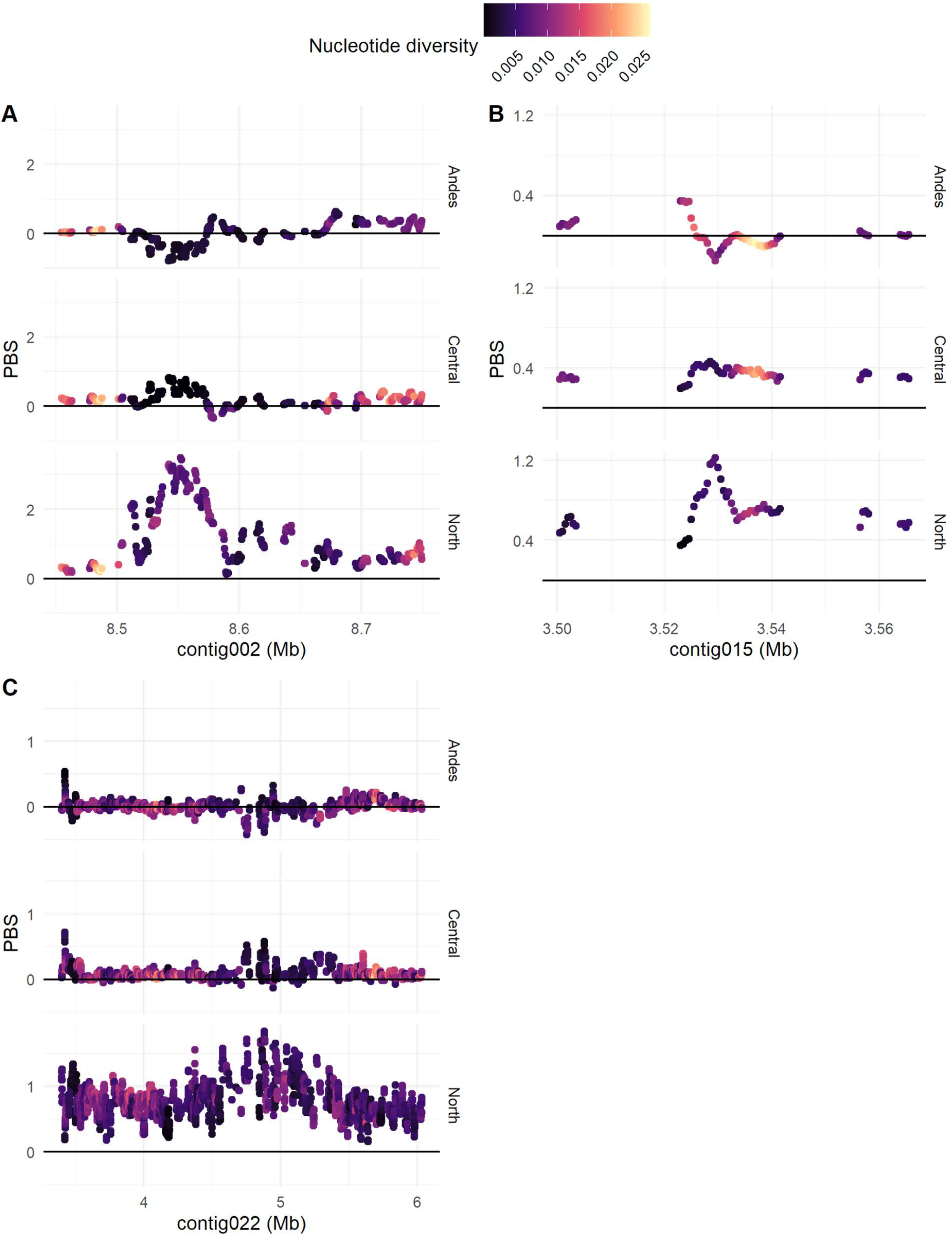
PBS and nucleotide diversity at three other PBS hotspots, with values averaged over 5kb intervals in 500bp steps.

## Supplemental Tables

**Table S1:** Sample names and collection locations.

| library | sample | site_abbreviation | sample_site | country |
| --- | --- | --- | --- | --- |
| C31 | Ind30 | AR | Arica-Arica State | Chile |
| C32 | Ind31 | AR | Arica-Arica State | Chile |
| C33 | Ind32 | AR | Arica-Arica State | Chile |
| C34 | Ind33 | AR | Arica-Arica State | Chile |
| C35 | Ind34 | AR | Arica-Arica State | Chile |
| C51 | Ind50 | CH | Chia-Cundinamarca State | Columbia |
| C52 | Ind51 | CH | Chia-Cundinamarca State | Columbia |
| C53 | Ind52 | CH | Chia-Cundinamarca State | Columbia |
| C54 | Ind53 | CH | Chia-Cundinamarca State | Columbia |
| C55 | Ind54 | CH | Chia-Cundinamarca State | Columbia |
| C11 | Ind10 | CN | Campo Nove-Caaguazu Department | Paraguay |
| C12 | Ind11 | CN | Campo Nove-Caaguazu Department | Paraguay |
| C13 | Ind12 | CN | Campo Nove-Caaguazu Department | Paraguay |
| C14 | Ind13 | CN | Campo Nove-Caaguazu Department | Paraguay |
| C15 | Ind14 | CN | Campo Nove-Caaguazu Department | Paraguay |
| C30 | Ind29 | CR | Costa Rica | Costa Rica |
| C56 | Ind55 | CR | Costa Rica | Costa Rica |
| C57 | Ind56 | CR | Costa Rica | Costa Rica |
| C58 | Ind57 | CR | Costa Rica | Costa Rica |
| C59 | Ind58 | CR | Costa Rica | Costa Rica |
| C41 | Ind40 | LC | La Curva-Arequipa Department | Peru |
| C42 | Ind41 | LC | La Curva-Arequipa Department | Peru |
| C43 | Ind42 | LC | La Curva-Arequipa Department | Peru |
| C44 | Ind43 | LC | La Curva-Arequipa Department | Peru |
| C45 | Ind44 | LC | La Curva-Arequipa Department | Peru |
| C36 | Ind35 | LJ | La Joia-Arequipa Department | Peru |
| C37 | Ind36 | LJ | La Joia-Arequipa Department | Peru |
| C38 | Ind37 | LJ | La Joia-Arequipa Department | Peru |
| C39 | Ind38 | LJ | La Joia-Arequipa Department | Peru |
| C40 | Ind39 | LJ | La Joia-Arequipa Department | Peru |
| C21 | Ind20 | MP | Mar del Plata-Buenos Aires State | Argentina |
| C22 | Ind21 | MP | Mar del Plata-Buenos Aires State | Argentina |
| C23 | Ind22 | MP | Mar del Plata-Buenos Aires State | Argentina |
| C24 | Ind23 | MP | Mar del Plata-Buenos Aires State | Argentina |
| C25 | Ind24 | MP | Mar del Plata-Buenos Aires State | Argentina |
| C01 | Ind0 | OV | Ouro Verde-Goias State | Brazil |
| C02 | Ind1 | OV | Ouro Verde-Goias State | Brazil |
| C03 | Ind2 | OV | Ouro Verde-Goias State | Brazil |
| C04 | Ind3 | OV | Ouro Verde-Goiás State | Brazil |
| C05 | Ind4 | OV | Ouro Verde-Goiás State | Brazil |
| PT1 | Ind67 | P.ope | Arvin, California | USA |
| PT2 | Ind68 | P.ope | Arvin, California | USA |
| PT3 | Ind69 | P.ope | Arvin, California | USA |
| PT4 | Ind70 | P.ope | Arvin, California | USA |
| PT5 | Ind71 | P.ope | Arvin, California | USA |
| PT6 | Ind72 | P.ope | Arvin, California | USA |
| PT7 | Ind73 | P.ope | Arvin, California | USA |
| PT8 | Ind74 | P.ope | Arvin, California | USA |
| C06 | Ind5 | RI | Riobamba-Chimborazo State | Ecuador |
| C07 | Ind6 | RI | Riobamba-Chimborazo State | Ecuador |
| C08 | Ind7 | RI | Riobamba-Chimborazo State | Ecuador |
| C09 | Ind8 | RI | Riobamba-Chimborazo State | Ecuador |
| C10 | Ind9 | RI | Riobamba-Chimborazo State | Ecuador |
| C46 | Ind45 | RO | Rocha-Rocha Department | Uruguay |
| C47 | Ind46 | RO | Rocha-Rocha Department | Uruguay |
| C48 | Ind47 | RO | Rocha-Rocha Department | Uruguay |
| C49 | Ind48 | RO | Rocha-Rocha Department | Uruguay |
| C50 | Ind49 | RO | Rocha-Rocha Department | Uruguay |
| C16 | Ind15 | SE | Santiago del Estero-Santiago del Estero State | Argentina |
| C17 | Ind16 | SE | Santiago del Estero-Santiago del Estero State | Argentina |
| C18 | Ind17 | SE | Santiago del Estero-Santiago del Estero State | Argentina |
| C19 | Ind18 | SE | Santiago del Estero-Santiago del Estero State | Argentina |
| C20 | Ind19 | SE | Santiago del Estero-Santiago del Estero State | Argentina |
| TA1 | Ind59 | SP | Barcelona, Catalonia | Spain |
| TA2 | Ind60 | SP | Barcelona, Catalonia | Spain |
| TA3 | Ind61 | SP | Barcelona, Catalonia | Spain |
| TA4 | Ind62 | SP | Barcelona, Catalonia | Spain |
| TA5 | Ind63 | SP | Barcelona, Catalonia | Spain |
| TA6 | Ind64 | SP | Barcelona, Catalonia | Spain |
| TA7 | Ind65 | SP | Barcelona, Catalonia | Spain |
| TA8 | Ind66 | SP | Barcelona, Catalonia | Spain |
| C26 | Ind25 | VA | Villa Alegre-Maule State | Chile |
| C27 | Ind26 | VA | Villa Alegre-Maule State | Chile |
| C28 | Ind27 | VA | Villa Alegre-Maule State | Chile |
| C29 | Ind28 | VA | Villa Alegre-Maule State | Chile |

**Table S2:** Maximum likelihood parameter estimates under the three population history models tested. See Figure S6 for model details.

| parameter | unit | M1 | M2 | M3 |
| --- | --- | --- | --- | --- |
| NANC | effective pop size | 3324287 | 3943689 | 3433648 |
| N0START | effective pop size | - | 187146 | 16394 |
| N1START | effective pop size | - | 121328 | 84783 |
| N0END | effective pop size | - | 2282860 | 1237043 |
| N1END | effective pop size | - | 2659059 | 857884 |
| N2END | effective pop size | - | 9698709 | 8346208 |
| T0 | generations ago | 523870 | 159990 | 187034 |
| T1 | generations ago | 1028654 | 239766 | 252383 |
| T2 | generations ago | - | 1082411 | 1134632 |
| T0EXPAND | generations ago | - | - | 184870 |
| T1EXPAND | generations ago | - | - | 230046 |
| R0 | exponential growth rate | - | -1.56E-05 | - |
| R1 | exponential growth rate | - | -1.29E-05 | - |
| R2 | exponential growth rate | - | -8.31E-07 | - |

**Table S3:** Allele frequency of mutations in either copy of ACHE1. Both A201S and F290V have been reported to confer organophosphate resistance.

|  | g8300 |  |  | g9292 |  |  |
| --- | --- | --- | --- | --- | --- | --- |
| mutation | North | Central | Andes | North | Central | Andes |
| A201S | 53% | 0% | 7% | - | 51% | 71% |
| F290V | 38% | 0% | 7% | - | 44% | 20% |
| F290N | 0% | 27% | 5% | - | 0% | 0% |

## Notes

### Competing Interest Statement

The authors have declared no competing interest.

